# Intracellular metabolic circuits shape inter-species microbial interactions

**DOI:** 10.1101/127332

**Authors:** Ali R. Zomorrodi, Daniel Segrè

## Abstract

Metabolite exchanges in microbial communities give rise to ecological interaction networks that influence ecosystem diversity and stability^1-6^. These exchanges depend on complex intracellular pathways thus raising the question of whether ecological interactions are inferable from genomes. We address these questions by integrating genome-scale models of metabolism^7^, to compute the fitness of interacting microbes, with evolutionary game theory, which uses these fitness values to infer evolutionarily stable interactions in multi-species microbial “games”. After validating our approach using data on sucrose hydrolysis by *S. cerevisiae*, we performed over 80,000 *in silico* experiments to evaluate the rise of unidirectional and cross-feeding metabolic dependencies in populations of *Escherichia coli* secreting 189 amino acid pairs. We found that, despite the diversity of exchanged amino acids, most pairs conform to general patterns of inter-species interactions. However, several amino acid pairs deviate from these patterns due to pleiotropy and epistasis in metabolic pathways. To better understand the emergence of cross-feeding, we performed *in silico* invasion experiments and found possible evolutionary paths that could lead to such association. Overall, our study provides mechanistic insights into the rise of evolutionarily stable interdependencies, with important implications for biomedicine and microbiome engineering ^8-10^.

---

Obligate dependencies among microorganisms, through the exchange of essential metabolites have been hypothesized to be ubiquitous in microbial ecosystems ^1^ Similar interactions have also been engineered in laboratory systems, mainly based on genetically induced auxotrophies ^3-6,11,12^. However, the evolutionary rise and maintenance of these interactions constitutes an unresolved puzzle, since genotypes that do not produce a given costly metabolite may have a selective advantage over producers. One theory, known as the Black Queen Hypothesis ^13^, suggests that metabolic dependencies could arise through adaptive gene loss: organisms that lose the capacity to produce a costly compound (non-producers) will have a selective advantage over organisms that produce and inevitably leak that compound (producers). This could give rise to an obligate dependency of non-producers on producers ^13^, or, in the case of more than one leaky function, to obligate cross-feeding (bidirectional dependency) ^14^. However, little is known about the conditions under which these dependencies would be established.

A limited number of theoretical studies have recently explored this question using ecological models ^15-18^, evolutionary game theory (see ^19-21^ for comprehensive reviews) and concepts from economics ^22^. While these approaches have provided valuable phenomenological insight into the general principles of metabolic interdependencies, they generally do not take into account the specific details of the organisms, pathways and molecules involved: behind the production, leakiness, and utilization of each metabolite, is a complex network of biochemical reactions, which may significantly vary from organism to organism, and across different conditions and exchanged metabolites. A powerful avenue to address this gap is the use of systems biology methods, such as stoichiometric models of metabolism ^7^. These models take into account the full metabolic circuitry of a cell and provide quantitative predictions of its growth capacity and metabolic fluxes. Recent work has started applying these approaches to model microbial communities (see ^23^ for a recent review) and to study the evolution of adaptive diversification in long-term evolutionary experiments ^24^. However, a systematic analysis of the underlying mechanisms and patterns of possible equilibrium states of inter-species interactions as a function of the leakiness of different metabolites is still unexplored. Here we propose a new hybrid modeling approach that combines the theoretical insight of evolutionary game theory with the organism-specific detailed analysis of cell-wide metabolic networks. We demonstrate how this strategy allows us to map the landscape of possible inter-species interactions, for which genome-scale models provide unique mechanistic insight.

Our mechanistic evolutionary game theory approach uses estimates of microbial fitness (payoff), based on genome-scale metabolic models, (forming the “payoff matrix” of the game). These payoffs are subsequently utilized to compute the Nash equilibria of “microbial interaction games” (see Methods). A Nash equilibrium is a central concept in game theory, defined as a state where no player can increase its payoff by a unilateral change of strategy. These payoffs also allows us to determine which of the identified Nash equilibria are evolutionarily stable by modeling the dynamics of genotype frequencies ^25^ (see Figure 1 and Methods). As a proof-of-concept of our approach, we modeled the yeast sucrose hydrolysis system^26^: when growing on sucrose, *S. cerevisiae* produces the surface enzyme invertase (encoded by the *suc2* gene), hydrolyzing sucrose into glucose and fructose, part of which serve as a public good. Since invertase production is metabolically costly, a mutant strain, which has lost its *suc2* gene, may emerge (see Figure 2A). Our simulations (based on the *iAZ*900 yeast metabolic model ^27^) reproduced the three types of equilibria observed experimentally ^26^, i.e. Prisoners’ Dilemma (non-producers dominate and the community collapses), Mutually Beneficial game (producers dominate), and Snowdrift game (producers and non-producers coexist) (see Figure 2B-2F, and supplementary text for details).

**Figure 1.**
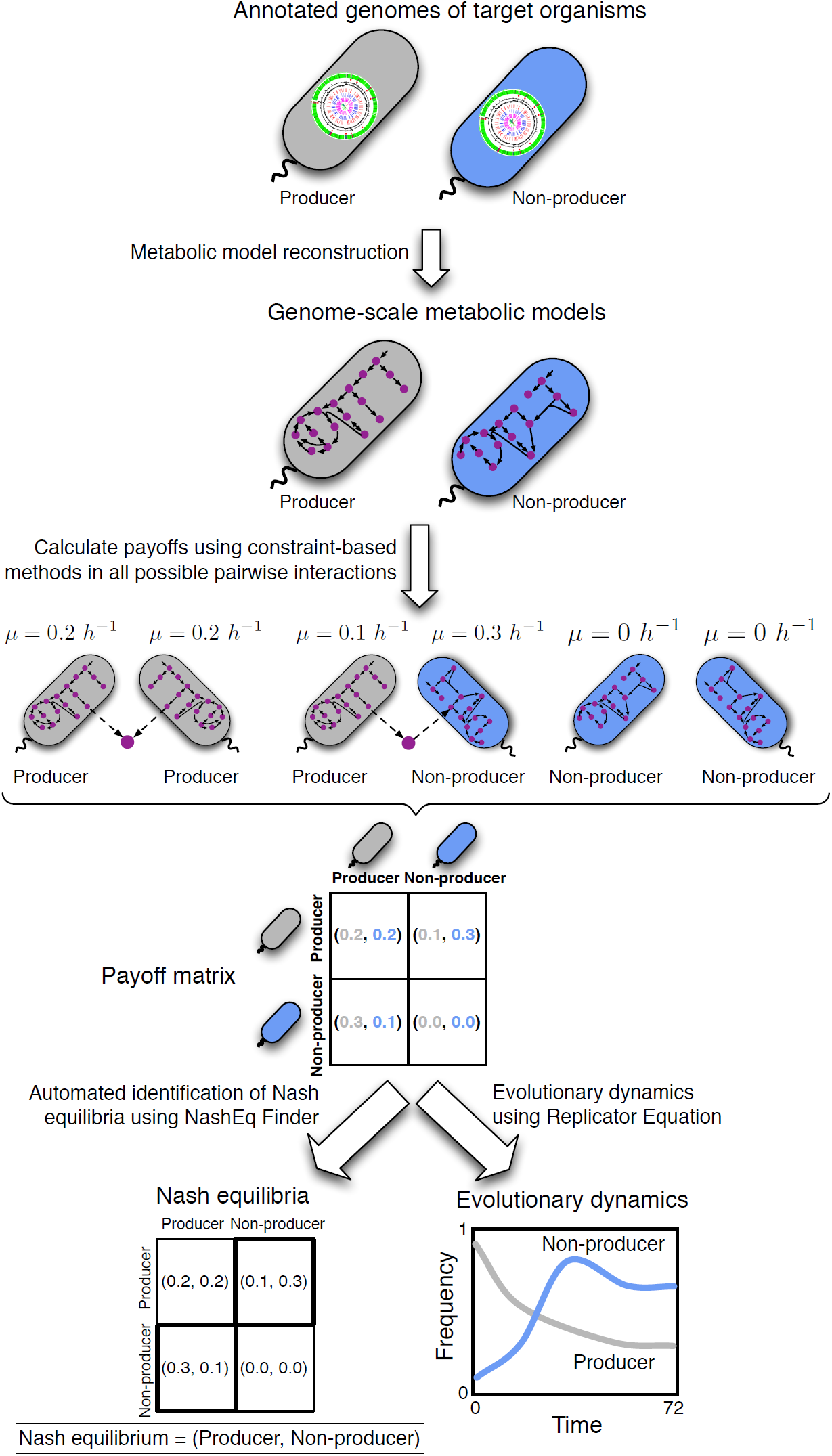
Outline of the proposed genome-driven evolutionary game theory approach. Annotated genomes of community members are used to construct genome-scale metabolic models. For each possible pair of genotypes in the community, constraint-based analysis tools for metabolic models such as Flux Balance Analysis ^34^ are used to estimate the fitness (or “payoff”) of each genotype as they engage in a specific metabolite-mediated interaction. These payoffs form the payoff matrix of the game. Based on this payoff matrix we identify all Nash equilibria of the game, using a newly developed automated tool (NashEq Finder, see supplementary text). The payoff matrix also allows us to model evolutionary dynamics (i.e., how genotype frequencies change over time) ^25^ and to determine which of the identified Nash equilibria are evolutionarily stable (see Methods).

**Figure 2.**
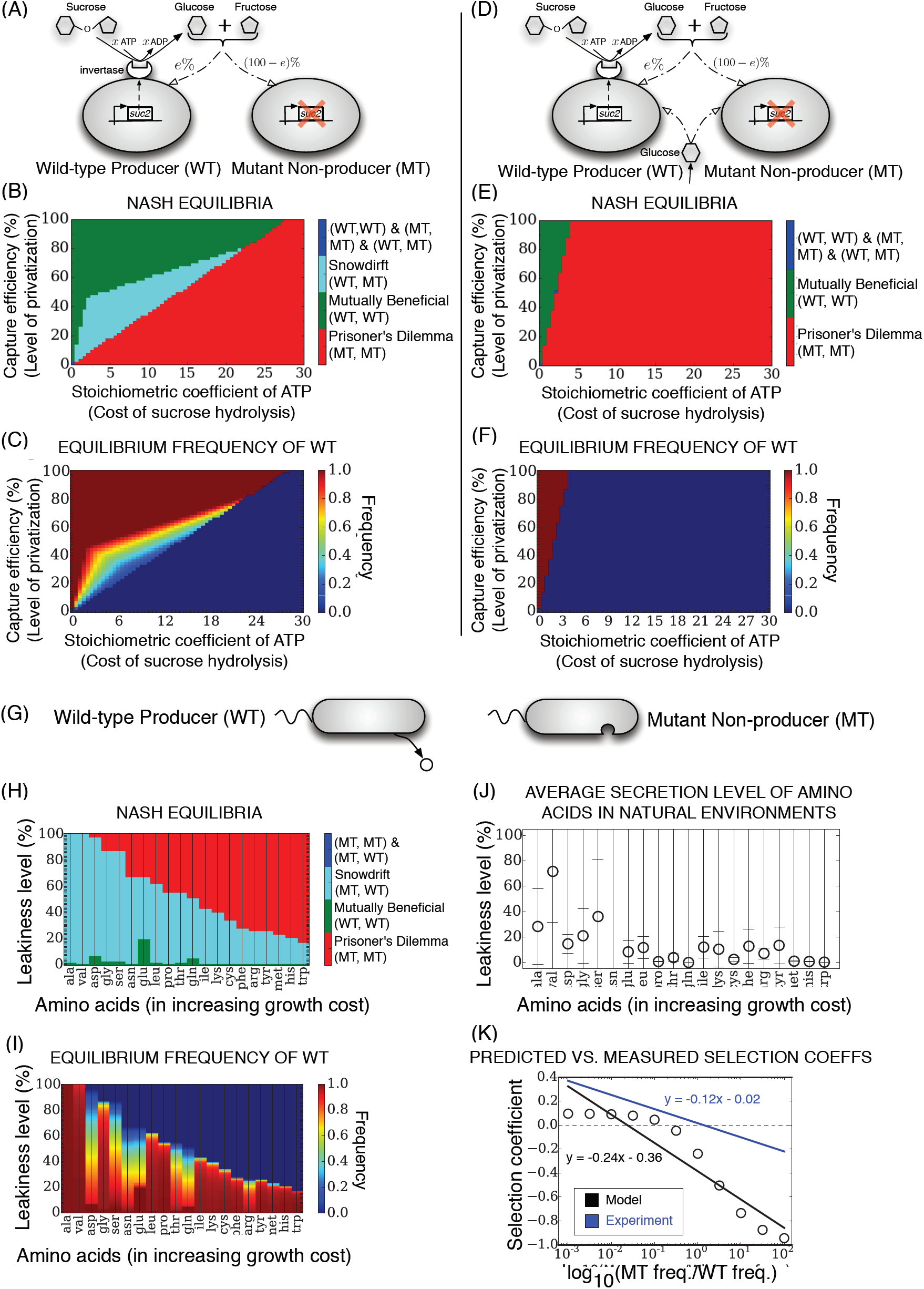
Metabolic dependencies in populations of *S. cerevisiae* growing on sucrose and in populations of *E. coli* secreting an amino acid. (A) Metabolic interactions between producer (wild-type, WT) and non-producer (mutant, MT) genotypes of *S. cerevisiae* growing on sucrose ^26^. Here, *e* represetns the percentage of glucose/fructose that diffuses away and serves as a public good. (B) Nash equilibria and (C) the equilibrium frequency of WT for the community shown in (A) as a function of the capture efficiency of the glucose/fructose and the invertase production cost (modeled by changing the stoichiometric coefficient of ATP in the sucrose hydrolysis reaction, indicated as *x*). An alternative *in silico* formulation of the energetic cost of invertase production that reproduces exactly the setup used in the experiment by Gore et al ^26^ (based on histidine auxotrophy) proved to be qualitatively equivalent to the analysis presented here (see supplementary text for details). The equilibrium frequency of WT was obtained from *in silico* invasion experiments (see Methods) for two cases of a small fraction of MT invading a resident population of WT and vice versa demonstrated that the equilibrium frequency of WT is the same in both cases. (D) Metabolic interactions between WT and MT when additional glucose is provided in the growth medium (see supplementary text for details). (E) Nash equilibria and (F) the equilibrium frequency of WT in the presence of glucose in the growth medium. The entire Snowdrift game region and part of the Mutually Beneficial region in (B) are replaced by the Prisoner’s Dilemma game, in (E), which is consistent with previous reports ^26^ and serves as an additional verification of our modeling approach. This is because in the presence of glucose, MT is less dependent on WT thereby increasing the average fitness of MT. (G) Possible genotypes in populations of *E. coli* leaking an amino acid include a prototrophic wild-type strain (WT) self-synthesizing a leaky amino acid and a mutant strain (MT) lacking the gene(s) for the biosynthesis of this amino acid. (H) The identified Nash equilibria for various leakiness levels (as a percentage of an *in silico* determined maximum: see supplementary text) across all 20 amino acids. Amino acids are shown here by using their standard three-letter code in the order of increasing *in silico* growth cost (see also supplementary Figure S1). (I) The equilibrium frequency of WT as a function of the leakiness level and amino acid type. *In silico* invasion experiments for two cases of MT invading WT and WT invading MT revealed that the equilibrium frequencies are insensitive to the initial frequencies. (J) Experimentally reported leakiness levels of amino acids averaged over three different datasets ^43,44^. Values in each dataset were normalized to their maximum (see supplementary Table S1 for values of data). Error bars show standard deviation over the three datasets. (K) Predicted selection coefficients across all amino acids and leakiness levels vs. the experimentally reported ones for *E. coli* ^29^. Empty circles show the average predicted selection coefficient for each amino acid. The black and blue lines show the fitted lines to the predicted and experimental values ^29^, respectively. The equation for the fitted line to experimental data was inferred from a graph of reference ^29^.

We next sought to characterize the landscape of possible ecological interactions in a large set of strains and exchanged metabolites. In particular, we explored the evolution of metabolic dependencies mediated by the leaky production of individual amino acids in *E. coli* and asked how the evolution of these dependencies varies across the 20 amino acids and different leakiness levels. Here, a prototrophic wild-type (WT, producer), leaking a given amino acid could interact with a mutant strain (MT, non-producer) lacking the gene(s) for the biosynthesis of that amino acid (see Figure 2G). The *i*JO1366 genome scale model of *E. coli*^28^ was used to construct *in silico* producer and non-producer strains, and to explore what equilibria can be established for a given leakiness level. As shown in Figure 2H, Prisoner’s Dilemma, Mutually Beneficial and Snowdrift outcomes are the major possible equilibria, similar to the yeast system (Figure 2B) (with the amino acid cost replacing the sucrose hydrolysis cost). However, a more complex pattern is observed here for the Mutually Beneficial region (see supplementary text for details), highlighting the organism- and product-specific nature of our model. In addition, one can observe that, for each amino acid, there is a threshold for leakiness level above which nonproducer mutants (MT) dominate, leading to community collapse (i.e., Prisoner’s Dilemma) (Figure 3H). Thus, we may expect the amino acids secretion levels in *E. coli* to lie below this threshold. Interestingly, the average of multiple published amino acid secretion datasets (see Figure 3J) displays a consistent trend of leakiness levels decreasing with increasing cost. To assess the evolutionary stability of above Nash equilibria we performed *in silico* invasion experiments, where a resident population of WT is invaded by a low frequency MT (and vice versa). This analysis showed that the equilibrium frequencies of WT and MT are independent of their initial frequencies (see Figure 3I). This had been theoretically ^14^ and experimentally ^14,29^ suggested to stem from the negative frequency dependence of fitness. In addition to recapitulating this pattern, our analysis provides a quantitative prediction of the selection coefficients of the 20 amino acids. As shown in Figure 3K, the predicted selection coefficients display reasonable agreement with previous experimental reports for *E. coli* ^29^. This establishes a nontrivial quantitative link between metabolic stoichiometry and important ecological parameters.

**Figure 3.**
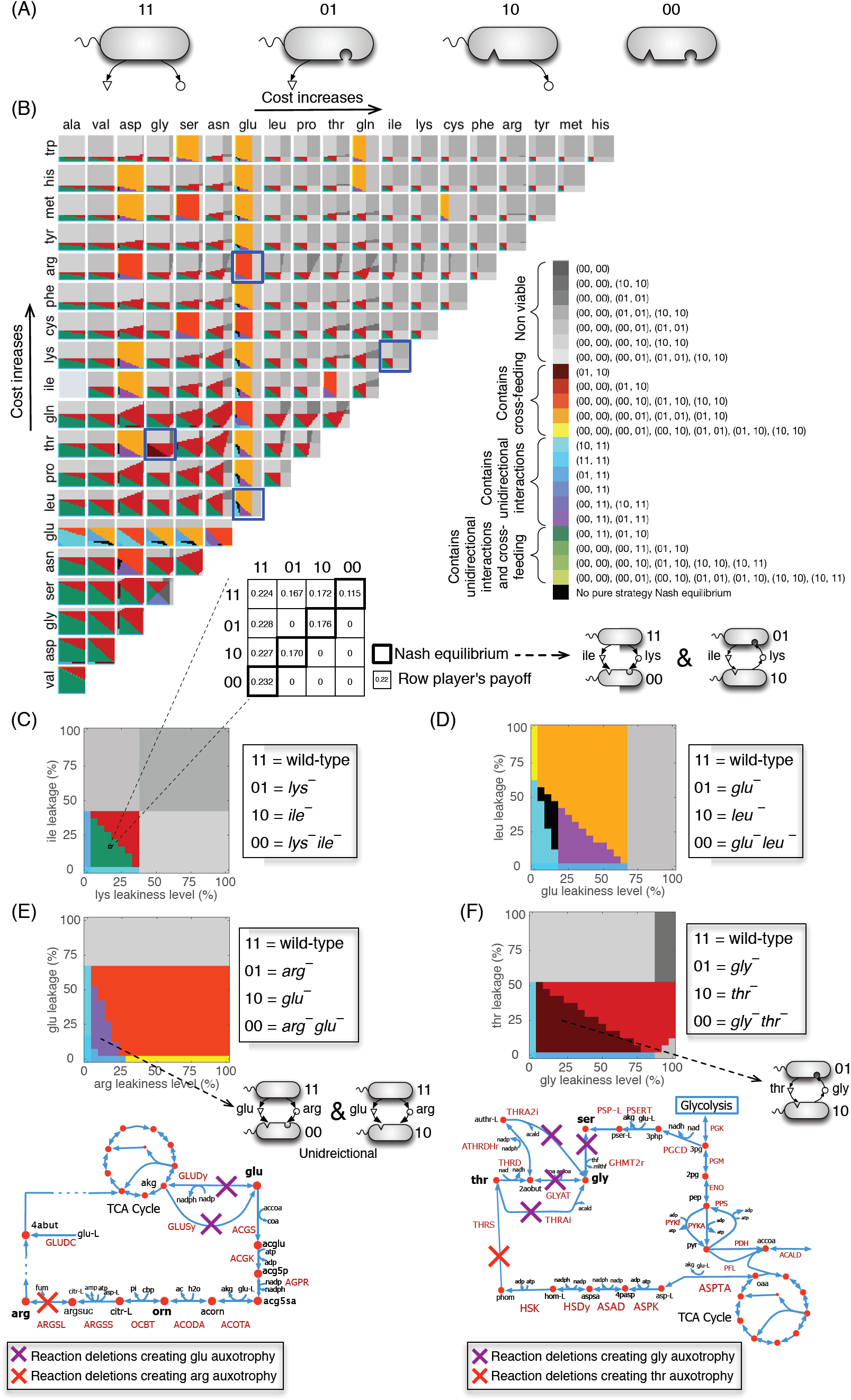
Equilibrium metabolic dependencies in populations of *E. coli* with two leaky amino acids. (A) Genotypes involved include a prototrophic strain self-synthesizing two leaky amino acids (i.e., 11), two single-mutant strains each lacking the gene(s) for the biosynthesis of one amino acid but synthesizing and leaking the other (i.e., 01 and 10), and a mutant strain lacking the genes for the biosynthesis of both leaky amino acids (i.e., 00). Here, ‘1’ and ‘0’ denote the presence or absence of biosynthesis pathways (genes) for an amino acid, respectively. (B) The identified Nash equilibria of two-player games (i.e., pairwise interactions) for all amino acids pair across different leakiness levels, zoomed in for four selected pairs including (C) (lysine, isoleucine), (D) (glutamate and leucine), (E) (arginine, glutamate) and (F) (glycine, threonine). A sample payoff matrix of the game is shown in (C) for a leakiness level of 15% for both lysine and isoleucine. Non-viable equilibria signify associations between genotypes that are non-viable leading to community collapse. Nash equilibria of three- and four-player games for a selected number of amino acid pairs are also given in supplementary Figures S2-S7. Metabolic maps in (E) And (F) show the metabolic pathways involved in the synthesis of the respective amino acids. Sample payoff matrices in the region of sustainable leakiness levels in (E) and (F) are given in supplementary Figure S8.

We further extended our analysis to map the landscape of ecological interactions in communities with two leaky amino acids. Under what conditions would the increased number of exchangeable metabolites give rise to more complex interspecies interdependencies, such as reciprocal exchanges? Four different genotypes are possible in this case (see Figure 3A): a prototrophic genotype that produces and leaks two amino acids (i.e., a full producer, denoted as ‘11’), two partial producer mutants each lacking the gene(s) for the biosynthesis of one amino acid (denoted as ‘01’ and ‘10’), and a no-producer mutant strain lacking the biosynthesis genes for both amino acids (denoted as ‘00’). In a community composed of all these four genotypes, different types of interactions are possible. Here we focus on pairwise interactions, such as cross-feeding, [01, 10] and unidirectional dependency, [00, 11], though higher-order interactions among three or four genotypes (e.g., [00, 01, 10] and [00, 01, 10, 11]) are possible as well. We next systematically computed all Nash equilibria, at varying leakiness levels, for 189 amino acid pairs, corresponding to all possible pairs of 20 amino acids (see Figure 3B) except for one, namely the (alanine, isoleucine) pair, because the 00 genotype for this pair is auxotrophic for a third amino acid, valine. (see supplementary Figures S2 to S7 for higher order interactions’ equilibria).

As shown in Figure 3B, a wide spectrum of equilibria ensues across different amino acid pairs and across different leakiness levels (e.g., see Figures 4C and 4D). Interestingly, despite the diversity of strains and exchanged metabolites, a majority of pairs are found to conform to general ecological patterns: for example, 139 (out of 189, or 73.5%) of amino acid pairs display a qualitatively identical region of leakiness levels (green regions in Figures 4B,C), where a unidirectional association, [00, 11], and cross-feeding, [01, 10], simultaneously emerge as Nash equilibria. The leakiness levels leading to this mixed equilibrium are the ones for which the full producer (i.e., 11 genotype) can still sustain growth. We refer to this region as *sustainable leakiness region* (see Methods). This is analogous to the maximum leakiness level in the study of individual amino acid secretions (Figure 2H): any leakiness levels for the two amino acids that lie outside this region would be expected to lead to extinction of the wild-type.

**Figure 4.**
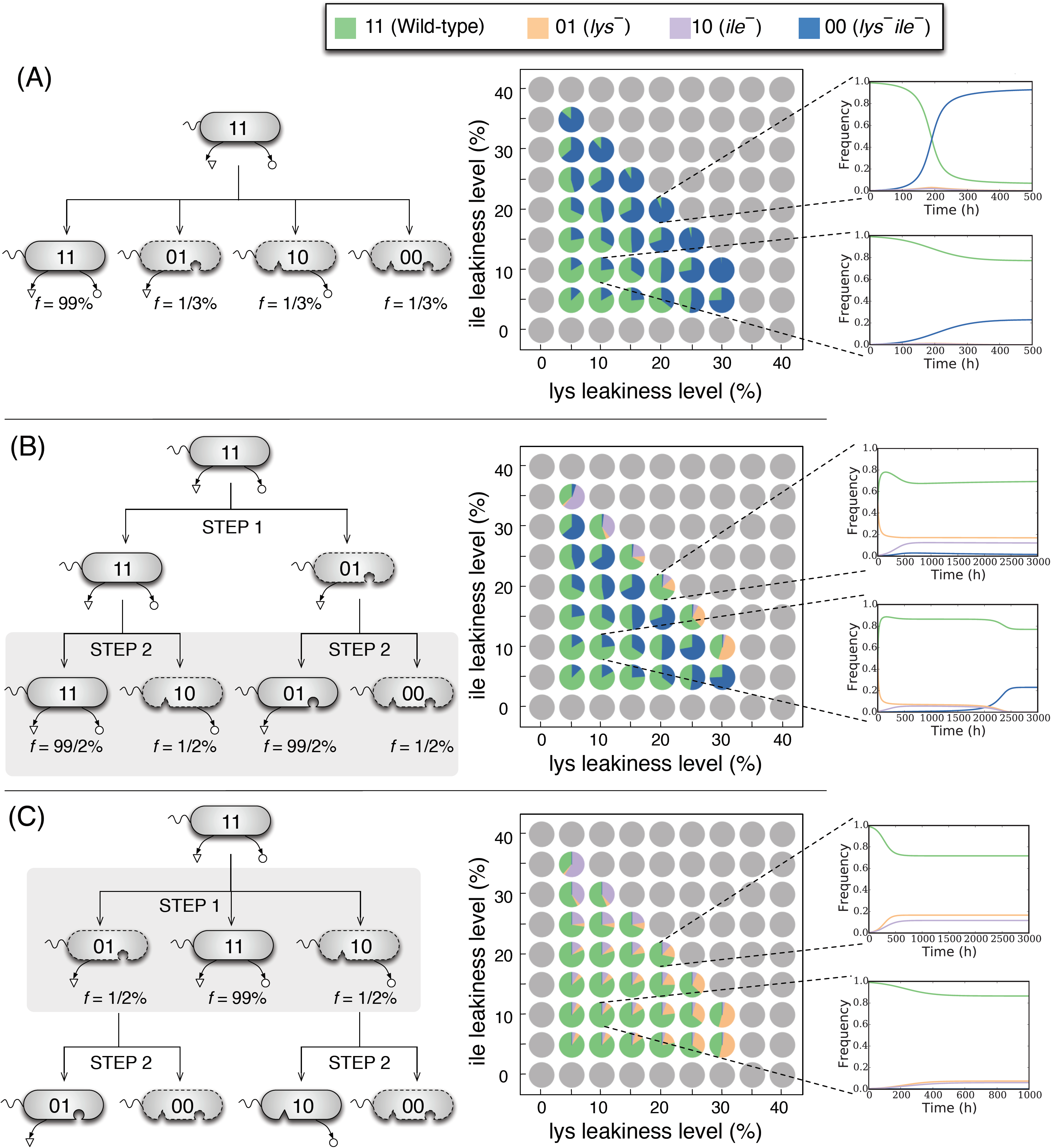
Impact of the initial genotype frequencies on the evolutionary emergence of metabolic dependencies in populations of *E. coli* with two leaky amino acids. Here, we have shown the results of targeted *in silico* invasion experiments for a representative amino acid pair (lysine, isoleucine) (see also Figure 3c). (A) 00, 01 and 10 simultaneously originate from 11 (i.e., wild-type) through genome streamlining and invade an existing population of 11 genotypes. (B) A small population of 10 and 00 invade a resident population of the 11 and 01. This simulates the second step of a two-step process for the loss of the leaky functions hypothesized in ^14^ (see the main text and also supplementary Figure S9). (C) A small population of 01 and 10 invades a resident population of 11. This models an alternative scenario for the two-step loss of leaky functions leading to stable cross-feeders: Two partial producer mutant genotypes (01 and 10) originate from 11, followed by the rise of the 00 genotype from 01 and/or 10. As shown here, cross-feeders can evolutionarily stabilize and co-exist with 11 genotypes in the first step. Further analysis showed that cross-feeders are also resistant to invasion by 00 genotypes arising in the second step (see supplementary Figure S10). Dynamic plots in (A)-(C) show the sample evolutionary dynamics of the system for selected equal leakiness levels for both lysine and isoleucine. Pie charts show the equilibrium frequencies of each genotype starting from the initial genotype frequencies shown in each panel. These equilibrium frequencies are given only for the sustainable leakiness region (green region in Figure 3C).

Another feature common to several amino acid pairs is the existence of a region in the leakiness plane (shown by red in Figure 3C) where cross-feeding (i.e., [01, 10]) is the only viable association, since excessive leakiness makes the full producer, 11, non-viable. This region is contiguous to the green region (with [00, 11] and [01, 10] as Nash equilibria). One interesting aspect of this configuration is that cross-feeding could initially ensue in the green region, coexisting with [00, 11] genotypes, and gradually move towards the red region (by evolving increased leakiness through selective advantage relative to the ancestor 11), leading to obligate cross-feeding as the only viable association.

In addition to exploring the landscape of Nash equilibria across different leakiness levels, it is interesting to ask whether the details of the biochemical networks underlying the genome-scale metabolic model predictions matter, and whether they provide direct explanatory power at the ecological level. Notably, for 50 amino acid pairs, the region of sustainable leakiness levels does not conform to the general pattern described above. For some pairs this region is partitioned into a number of sub-regions each corresponding to a different equilibrium (e.g. see Figure 3D) while in some extreme cases this entire region corresponds to a single Nash equilibrium, e.g., for (arginine, glutamate) and (glycine, threonine) (see below).

For the first anomalous pair, (arginine, glutamate), Nash equilibria in the sustainable leakiness region includes [10, 11] and [00, 11], but not the cross-feeding state [01, 10] (see Figure 3E). Inspection of the biosynthesis pathway of arginine and glutamate in *E. coli* revealed that glutamate is required for the production of ornithine, which serves as an essential precursor for the biosynthesis of arginine. This implies that a mutant strain lacking the biosynthesis pathways for glutamate (i.e., strain 10) will not be able to synthesize and leak arginine, thus acting like a 00 genotype and preventing the occurrence of cross-feeding. This observation is consistent with a previous report on the inability of arginine and glutamate auxotrophic mutant strains to grow in a co-culture under the minimal medium ^3^. This effect is due to a pleiotropic metabolic gene (i.e. a gene whose modification affects more than one metabolic phenotype), and illustrates how core biochemistry can impact ecological interactions.

A more complex scenario occurs for the (glycine, threonine) pair, where cross-feeding, [01, 10], is the only Nash equilibrium that emerges in the sustainable leakiness region (see Figure 3F). What prevents [00, 11] from being a Nash equilibrium? One of the conditions for [00, 11] to be a Nash equilibrium is that, in presence of 11, the fitness of the non-producer (00) should be higher than any of the partial producers (01 and 10) (as one would intuitively expect, because 00 does not incur the production cost of the two amino acids). In this case, however, it turns out that 00 is less fit than 10 (*thr*^−^ mutant, see a sample payoff matrix in supplementary Figure S8) thereby preventing [00, 11] from being a Nash equilibrium. This anomalous effect is due to the fact that the concurrent removal of glycine and threonine biosynthesis genes (in 00) will lead to a reduction in the capacity to produce other essential biomass components (such as serine). Interestingly, this is a case of diminishing-return (or “negative”) epistasis, in which the effect of the double mutation (00) is less severe than expected based on the two single mutations (01 and 10). Negative epistatic interactions preventing the appearance of [00, 11] as the Nash equilibrium can be observed for 34 other amino acid pairs (see supplementary Table S1). This pattern is supported by existing experimental reports showing that epistasis correlates negatively with the expected fitness of multiple “genome streamlining” mutations in *E. coli*, thereby causing diminishing returns ^30^. These results provide mechanistic insights into how epistatic interactions among intracellular pathways can affect ecological interactions, a feature that cannot be captured by abstract phenomenological models.

It is next interesting to ask whether our approach can shed light onto the possible paths towards the rise of different interactions, especially cross-feeding. In the landscapes of identified Nash equilibria (Figure 3B), cross feeding ([01, 10]) emerges together with other equilibria (such as [00, 11], e.g. in the green region in Figure 3C). This raises the question of whether and under what conditions cross-feeding would be evolutionarily stable. By performing a number of targeted *in silico* invasion experiments (see Methods), we found that this depends strongly on the initial frequencies of the different genotypes (shown in Figure 3A). In particular, our analysis demonstrates that cross-feeders (01 and 10) will go extinct if they invade the full producer (11) in presence of the non-producer genotype (00) (see Figure 4A). However, the cross-feeders can subsist if 00 invades at a later stage, e.g. if it arises from mutations in the partial producers (01 or 10) (see Figures 4B and 4C for two such mechanisms). An example of the latter scenario is the two-step process hypothesized in ^14^: first, the biosynthetic capacity for one amino acid is lost, e.g. resulting in a 01 genotype, which could equilibrate and co-exist with 11 (see Figures 2G-2I). In the second step, either 01 or 11 may lose their capacity for producing the second amino acid (because the other strain can compensate), giving rise to 00 and 10 genotypes, respectively (see Figure 4B). Here, we quantitatively explored this hypothesis by assessing the evolutionary dynamics for the second step, assuming that equilibration of the first step has already occurred. As shown in Figure 4B, the prototrophic (11) and no-producer (00) genotypes always survive in this competition, while cross-feeders evolutionarily emerge only at high leakiness levels. Further *in silico* invasion experiments demonstrate that established cross-feeding pairs tend to be resistant to invasion by non-producers (see supplementary Figure S10) and by prototrophs (see supplementary Figure S11). Thus, once established, obligate mutual metabolic exchange could be evolutionarily stable against invasion by other genotypes, even in a homogenous environment, consistent with previous experimental reports ^31^. Notably this stability against invasions is dependent on the specific metabolites exchanged and on the level of metabolic exchange (leakiness).

We demonstrated here that by adding new layers of details to abstract theoretical ecology models, we can reveal how intracellular molecular mechanisms (including pleiotropy and epistasis of metabolic enzyme genes) lead to the rise of ecological interactions. The analysis we presented spans over 80,000 *in silico* experiments (across 189 amino acid pairs and 441 leakiness level combinations), which is beyond the current experimental capabilities. This study provides a guideline for the design of future targeted experiments, e.g. built upon previously established synthetic communities ^3-6,11^. For example, our computational results could be used to suggest choices of metabolite pairs and ranges of engineered leakiness levels that lead to the establishment of a specific inter-dependency, e.g., unidirectional vs., cross-feeding. In addition, our study offers a basis for better understanding of metabolic interdependencies in natural microbial communities, such as those in the human gut microbiota ^8-10^. From the perspective of biotechnological applications, our approach lays the foundation for proactively incorporating evolutionary concepts in the *de novo* design of synthetic microbial consortia that are resistant to invasion by competing strategies ^23^.

## Methods

### Background on evolutionary game theory

Evolutionary game theory is the application of classical game theory to model the evolutionary dynamics of mixed populations. Modeling microbial communities with evolutionary game theory involves two steps: (i) Considering all pairwise interactions among genotypes and estimating the payoff (fitness) of a microbial player *k* upon interacting with a partner *k*′. Higher order interactions can be similarly defined; These estimated payoffs are represented in the form of a matrix, referred to as payoff matrix. (ii) Using the payoff matrix to identify the Nash equilibrium(s) - a fundamental concept in game theory defined as a state where no player has an incentive to unilaterally change its current strategy, because it cannot improve its payoff by doing so. An *evolutionary stable strategy* is a similar concept in evolutionary game theory: It is a Nash equilibrium, which is evolutionarily stable, i.e., natural selection alone is sufficient to prevent invasion by competing mutant strategies. Evolutionary stable strategies can be found by modeling the evolutionary dynamics of the game using the computed payoffs (see the following sections). In this text, by ‘evolutionary dynamics’ we mean how the relative genotype abundances (frequencies, or community structure) change over time following evolutionary game theory ^25^, and microbial ecology literature ^32^. This aspect of evolutionary dynamics, as opposed to the simulation of *de novo* mutations and subsequent selection processes (which have been pursued in other studies ^33^), is the main focus of our analysis.

### Background on Flux Balance Analysis (FBA)

FBA (described in detail elsewhere, e.g. ^34^) is a linear optimization problem that uses genome-scale metabolic models to make quantitative predictions about the cell’s growth rate, intracellular reaction fluxes and secretion rates of metabolites that are potentially excreted by the cell under a given condition. In the most common formulation, this is achieved by maximizing the flux of a pseudo-reaction called *biomass reaction* (*v*_*biomass*_) whose reactants are precursors required for growth and whose flux is indicative of the cell’s growth capacity. This is subject to constraints imposing the steady-state mass balance for each metabolite in the network (see Constraint 1 below) and lower and upper bounds on reactions fluxes to impose the reversibility of reactions and uptake and aeration conditions (see Constraint 2 below). The standard formulation of FBA is as follows:

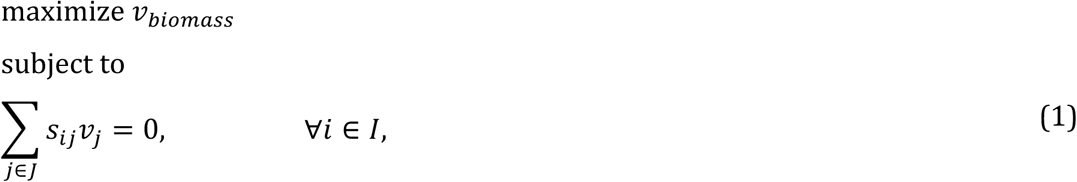

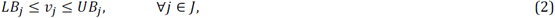
where, *I* is the set of metabolites, *J* is the set of reactions, *s*_*Ij*_ is the stoichiometric coefficient of metabolite *i* in reaction *j* (known from the metabolic model), *LB*_*j*_ and *UB*_*j*_ denote lower and upper bounds on flux of a reaction *j* (provided as inputs), respectively, and *v*_*j*_ is the flux of a reaction *j* (optimization variables).

### Using genome-scale models and FBA to compute payoffs for interacting microbes

We used FBA to provide organism-specific and genomically-informed estimates of the payoffs upon specific pairwise interactions (see also Figure 1). For a given pair of genotypes *k* and *k*′, we solve a separate FBA problem for *k* and *k*′. Each FBA problem involves two new types of constraints added to the standard FBA: (i) The first type of constraints pertains to implementing *in silico* the gene deletions that correspond to the genotype under consideration (see Equation 3 below). For example, if genotype *k* is auxotroph for lysine, this auxotrophy can be in principle induced by knocking out gene *lysA*, which codes for diaminopimelate decarboxylase. (DAPC). In the FBA model, this gene deletion is simulated by setting *v*_*DAPDC*_ = 0. More complex gene-to-reaction mappings could lead to more complex set of constraints. In cases where several different choices are possible for genes whose deletion would induce a given auxotrophy, we select one. (ii) The second type of constraints (Equations 4 and 5) simulates the exchange of metabolites between the genotype under consideration and its partner(s). In our calculations, for any given exchanged metabolite (e.g., amino acid or amino acids pair), we systematically explore the effects of a range of possible leakiness levels. Thus in each *in silico* experiment, the leakiness levels of amino acids secreted by each genotype are fixed at a pre-specified value. Similarly, metabolites leaked by the partner are made available to this genotype through appropriate upper bounds on import fluxes. The optimal value of biomass flux obtained upon solving this FBA problem provides an estimate of the growth rate of each genotype in a given pairwise interaction, which we use as a proxy for its payoff. For example, the payoff of *k* when facing *k*′ (*a*_*kk*′_) is 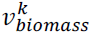 and the payoff of *k*′ when facing *k* (*a*_*k*′ *k*_) is 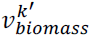. The general form of the FBA formulation for each genotype is mathematically formulated as follows:

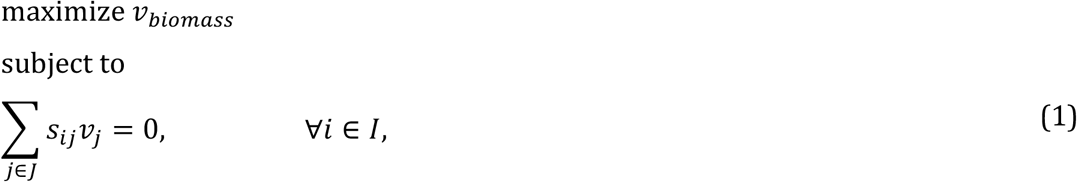

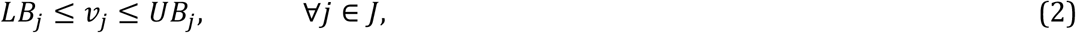

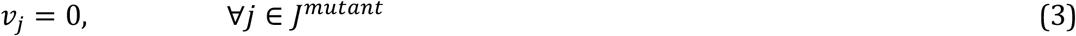

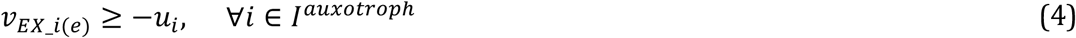

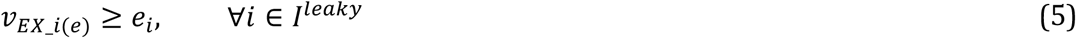
where, *J*^*mutation*^ denotes the set of reactions corresponding to specific gene mutations for the genotype under consideration. In addition, *I*^*auxotroph*^ ⊂ *I* represents the set of metabolites that this genotype is auxotroph for, but that are provided by other genotypes, and *I*^*leaky*^ ⊂ *I* is the set of leaky (secreted) metabolites by the genotype under consideration. *v*_*EX_i(e)*_ denotes the flux of exchange reaction for a metabolite *i. u*_*i*_ > 0 and *e*_*i*_ > 0 denote the pre-specified net uptake and export flux of a metabolite *i*, respectively. Input parameters for this FBA problem include: (*i*) The list of metabolites that each genotype is leaking and those available for uptake from the partner genotype in a pairwise (or higher order) interaction, (*ii*) The leakiness level of exchanged metabolites, and (*iii*) the net uptake and export fluxes of exchanged metabolites (i.e., *e*_*i*_ and *u*_*I*_) calculated based on the fixed leakiness levels and the specific interacting genotypes (see supplementary text for details). Constraint (3) sets to zero the flux of reactions corresponding to the specific gene mutations in the genotype under consideration. Constraints (4) allows for the uptake of metabolites available from partner genotype(s) in a pairwise or (higher-order) interaction. Constraint (5) requires the export of leaky metabolites at the pre-specified level *e*_*i*_. The payoff of the genotype under consideration is set to the optimal value of the biomass flux, or to the death rate (a negative value), in the case of an infeasible problem. An infeasible FBA problem may occur due the lack of enough carbon source to satisfy maintenance ATP requirements in the model or due to imposing a high level of leakiness for a leaked metabolite. Imposed leakiness level causing this infeasibility are referred to ‘unsustainable leakiness levels’. The details of specific formulations for the presented case studies with *S. cerevisiae* and *E. coli* are given in the supplementary text. Additional environmental/strategic/genetic conditions can be incorporated through the addition of appropriately defined constraints. In addition, one can use a different objective function (e.g., the minimization of metabolic adjustment ^35^) or other constraint-based community modeling tools e.g., ^36,37^ as an alternative.

### Optimization-driven automated identification of the Nash equilibria

Upon constructing the payoff matrix, as described above, one can identify the Nash equilibrium(s) of the game. We developed an efficient optimization-based procedure called *NashEq Finder* to automate the identification of all pure strategy Nash equilibria of a game given its payoff matrix. NashEq Finder is an integer linear program (ILP), which relies on binary variables to decide on whether or not a particular entry of the payoff matrix satisfies the conditions of a Nash equilibrium (see supplementary text for a detailed formulation). This algorithm is able to identify all possible Nash equilibria of a game with any number of players and thus can be reliably used for the rapid identification of the equilibrium states of complex communities.

### Modeling the evolutionary dynamics at a genome-scale resolution

Following standard approaches in evolutionary game theory we model evolutionary dynamics using the Replicator Equation ^25^. In particular, to take into account interactions higher than pairwise, we used an extended form of the classical Replicator equation^25^. This equation predicts the changes in the relative abundance (frequencies) of genotypes over time according to their reproductive fitness under the assumption of a roughly constant population size ^25^ (see supplementary text for the difference between this equation and multi-species dynamics flux balance analysis ^38^). Our formulation, following ^39^, can be expressed as follows:

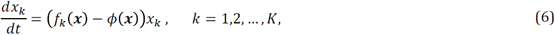

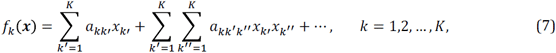

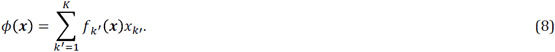

Here, *K* denotes the set of community members and ***x*** = [*x*_1_, *x*_2_, …, *x*_*K*_]^*T*^ is the composition of the community with *x*_*k*_ being the frequency of genotype *k*. *f*_*k*_(***x***) is the average reproductive fitness of genotype *k* that depends not just on other genotypes it may encounter but also on their frequencies. *a*_*kk*′_ and *a*_*kk*′ *k*″_ denote the payoffs of genotype *k* when encountering another genotype *k*′ in a two-player game (i.e. pairwise interaction) or two other genotypes *k*′ and *k*″ in a three-player game, respectively. Finally, *ϕ*(***x***) is the average fitness of the entire community.

Here we use the replicator equation to perform targeted *in silico* invasion experiments, in which a newly emerged low frequency genotype invades an existing resident genotype. While such *in silico* experiments are based only on evolutionary dynamics (modeled through the Replicator equation), future developments could take into account additional aspects such as eco-evolutionary feedback ^32,40-42^.

